# Cryptic speciation in freshwaters: are speciation in lentic species shaped by paleohydrological events?

**DOI:** 10.1101/2021.05.29.446258

**Authors:** Maitreya Sil, Abhishikta Roy, Tenzin Palden, K. Praveen Karanth, N. A. Aravind

## Abstract

The Indian subcontinent is extremely diverse in terms of its flora and fauna. However, there is a severe underestimation of the biotic diversity in invertebrate groups as well as a lack of understanding of the processes generating diversity in these groups. Here we aimed to jointly estimate the cryptic diversity of two freshwater snail species complexes – Pila globosa and Pila virens, and uncover the processes behind the speciation in these groups. We employed phylogenetic, morphometric, population genetic and niche modelling technique to address these questions. We found out that both species complexes consist of several genetically and ecologically distinct putative species. The speciation was primarily driven by allopatric isolation into different river basins. A combination of paleoclimatic and paleohydrological changes during the Miocene have shaped the speciation events. Local climatic adaptation also could have contributed towards some of the speciation events. The study sheds light on the complex interaction between the biology of the species and the environment that shape the diversification patterns in a group.

## Introduction

Species are fundamental units of macroevolution, as well as of several ecological concepts. Hence, it is imperative to estimate the number of species belonging to a lineage or distributed in an area to understand the evolutionary and ecological processes behind the diversity. None of the existing seem to unanimously fulfil all the criteria required for delineating species. (Queiroz, 1998) attempted to resolve this issue by classifying species as separately evolving populations and categorizing all the fundamental concepts of the existing species concepts as species criteria, all or some of which arise at different levels of lineage divergence. Thus, any of these criteria are enough to delineate species, however, multiple lines of evidence confer more confidence to the degree of species divergence. Morphologically cryptic species pose a hurdle for morphology-based taxonomy. Integrative taxonomic framework which uses multiple lines of evidence in compliance to unifying species concept, has been successfully implemented in resolving species complexes consisting of multiple cryptic species (Karanth, 2017). An understanding of the diversity is intricately linked to the processes that generate diversity (Wiens, 2004). Speciation can result from geographic isolation or ecological divergence in the absence of geographic separation. Often geographic isolation leads to subsequent ecological divergence. It is imperative to understand the interplay between geography and ecology to gain foresight into the speciation process.

Indian subcontinent provides an ideal setting to address various questions pertaining to species diversity and speciation owing to its geographic and climatic attributes (Mani, 1974; Valdiya, 2010). Although part of Asia, the subcontinent is functionally insular owing to presence of geological or climatic barrier to dispersal into or out of India. This has led to the evolution of several in-situ diversifications, similar to other insular systems such as islands, lakes etc. (Praveen Karanth, 2015). Several studies have shown presence of cryptic lineages in the subcontinent in a range of taxonomic groups, such as langurs, Sitanas (Deepak and Karanth, 2017), ground-dwelling geckos (Lajmi et al., 2016), centipedes (Joshi and Karanth, 2012) etc. The subcontinent exhibits extreme geographic heterogeneity, which might serve as potential driver of speciation (Vijayakumar et al., n.d.). The subcontinent also underwent extreme climatic changes from being warm and humid in Paleocene to being progressively dry and seasonal (Clift et al., 2008; Licht et al., 2014; Molnar et al., 1993; Nelson, 2007; Zachos et al., 2001). The climatic oscillation has catalysed diversification in several lineages (Agarwal and Ramakrishnan, 2017; Deepak and Karanth, 2017; Lajmi and Karanth, 2020). However, speciation in freshwater groups is rarely explored in the Indian context.

Freshwater snails are an understudied group that needs major taxonomic revisions in order to address pertinent evolutionary and ecological questions, assess their conservation status, and assess their importance as bioresources. One such group is the genus *Pila*, distributed in Africa and Asia. Prashad noted down six species of *Pila* from India (Prashad, 1924). It has been observed that often widespread species tend to be species complexes consisting of multiple cryptic species. Two of these species *P. globosa* and *P. virens* are widespread. The species P. globosa is distributed in current day Maharashtra, Central India, Rajasthan, Throughout the Gangetic plains from Uttarakhand to West Bengal, Orissa and parts of Northeast India. The species might also be found in Sri Lanka. Prashad reported the species *P. virens* from Maharashtra, Tamil Nadu and Assam, although more recent literature suggested more widespread distribution in Peninsular India and does not report any sightings from Assam. In addition to being widespread, these species show high intra-species divergence, suggesting that they might actually be species complexes.

Freshwater snails are also excellent model systems to understand speciation. On one hand there are instances of extensive radiations within lake systems through trophic or habitat specialization (Sengupta et al., 2009). There are also examples of allopatric speciation in rives and other freshwater systems. Allopatric speciation in freshwater systems are often associated with paleohydrological changes such as river capture events or marine incursions (Lundberg et al., 1998). However, dispersal in freshwater snails across river basins are also known to be aided by aquatic birds (Kappes and Haase, 2012). Finally, *Pila* is particularly interesting, since, it is amphibious and can disperse on land over short distances. It is imperative to investigate the role played by river systems as well as ecology in driving speciation in different species of Pila species complex. Since, these species occupy multiple river basins across the Indian subcontinent, it is likely that river basin segregation-driven allopatric species was the primary cause of speciation in this group as observed in various fish genera (Lundberg et al., 1998). On the contrary the speciation events could have been sympatric taking place in the same river basin. Sympatric speciation has been observed in Pachychilid snails in Kaek river system in Thailand (Glaubrecht and Köhler, 2004), and in the Poso and the Malili lake systems of Sulawesi (Matthias Glaubrecht, Von Rintelen, 2008) in *Lanistes* (Schultheiss et al., 2009) and *Bellamya* (Schultheiß et al., 2011) in great lakes of Africa. The extreme climatic variation in the subcontinent could have contributed as well. Finally, climatic shifts are known drivers of speciation. Historic climatic oscillation in the Indian subcontinent since Paleocene also could have contributed to the speciation event. A thorough understanding of the distribution, ecology and evolutionary history is necessary to uncover these processes.

Here, we implemented an integrative taxonomic approach combining molecular, morphometric and species distribution modelling tools to uncover the diversity of each of the described widespread *Pila* species from the subcontinent. We addressed the following questions: 1) Are the described species of *Pila*, in reality species complexes consisting of more than one species? 2) were these speciation events allopatric or sympatric? 3) What role did paleohydrological events or ecology play in driving speciation in these species complex?

## Methods

### Sample Collection

Whole specimens were collected from all over the distribution range of each widespread species, as well as from and around the type localities of the data deficient species. Samples were preserved in absolute ethanol. The latitude and longitudes of the collection localities were noted down in order to carry out species distribution modelling (Table S1).

### Gene Sampling

Genomic DNA was extracted from the foot muscle tissue of individual specimens using the CTAB extraction method (Chakraborty et al., 2020). A fragment of the COI gene was amplified for each individual. One, species delimitation analyses were carried out, fragments of the 18s rRNA and Histone H3 gene were also PCR amplified for a few individuals from those putative species where nuclear data was not available (Table S2). The amplicons were sequenced in Barcode Biosciences.

### Phylogenetic and Species Delimitation analyses

The obtained COI sequences were aligned in Mega7 (Kumar et al., 2016). The alignment consisted of the data generated for the current study, as well as *Pila* sequences from India, Southeast Asia and Africa. Two African genera, *Lanistes* and *Afropomus* were used as outgroups. The models of sequence evolution and per codon partition scheme was established using PartitionFinder v2 (Lanfear et al., 2017). Bayesian inference phylogeny was reconstructed using MrBayes v3 (Ronquist and Huelsenbeck, 2003). Two parallel runs were performed, and the saturation of the runs were estimated through observation of the lowering of standard deviation of split frequency till below 0.01. Furthermore, the parameter files were imported into Tracer v1.7 to check thorough sampling of the tree space (ESS>200). The BI trees obtained were subjected to mPTP species delimitation analyses (Kapli et al., 2017). The mPTP analyses were carried out in the mPTP server and all the parameters were kept at default.

### Morphometric analyses

Morphometric measurements were carried out on select individuals from each putative species suggested by mPTP. A Vernier callipers was used to take measurements. Nine shell characteristics were measured. Ratios of informative characters were computed from these initial characters in order to size correct these measurements (Table S3). The correlated variables were removed from further analyses. Next, we carried out PCA to observe if different putative species can be distinguished on the PC axes based on the measurements. Furthermore, ANOVA was carried out based on the ratios that contributed the most to PC1 to ascertain whether the differences between species were significant. All the above-mentioned analyses were carried out in R. In order to understand if the morphometric data can be separated into clusters, we carried out partitioning around metoids (PAM) method. The PAM method assigns data points to the closest among k number of metoids. First optimum number of clusters were calculated from the ratios, by comparing the silhouette width of clusters ranging from two to ten. Next, the final analyses were carried out after specifying the optimum number of clusters. The analysis was carried out in R using the package Cluster. All the above mentioned analyses were carried out separately for *P. globosa* and *P. virens* complex.

### Geometric morphometry

Geometric morphometry is a digital landmark-based approach. The specimens were photographed using a Nikkon P900 camera, against a white background. The JPEG files were converted to one single TPS file using tpsUtil32 v1.7.4. Thereafter the TPS file was then imported into tpsdig232. A total of 19 landmarks were placed in order to capture the variation of the shell shape between putative species. The final analyses was carried out in R using the package Geomorph. Out of the 19 landmarks, 15 were treated as sliding semilandmarks. First a Generalized Procrustes Fit was performed in order to transform the shape spaces into tangential space, aligned by the principal axis, Next, PCA analyses was performed from the coordinates of the Procrustes transformed data to visualize whether the putative species can be differentiated based on shape. Finally, a permutation-based distance was computed between groups to observe if the groups are significantly different from one another.

### Species Distribution Modelling

Species distribution modelling was carried out using MaxEnt Ver 3.4k (See below). The nineteen bioclimatic variables (Bio 1 to Bio 19; See Supplementary Table 1) with the spatial resolution of 30 arc seconds (approximately 1km2) for current climate were downloaded from WorldClim Database version 2 (Hijmans et al., 2005) (http://www.worldclim.org/). The potential geographic distribution of *Pila* spp. was modelled using Maximum entropy algorithm model (MaxEnt) version 3.4.1k (Philips et al., 2006) for the current climate scenario. MaxEnt approximations are based on the probable occurrence of species records finding the maximum entropy distribution with high extrapolative accuracy. To avoid model complexity and multi-collinearity, only xxx to yyy variables (Supplementary Table 1) were retained for further analysis based on their low degree of correlation between themselves (Pearson’s coefficient value of |r| <= 0.7). For modelling we used randomly selected 25% as test data and the remaining 75% was used for model training. We used the default feature in MaxEnt to run the model. The area under the receiving operator curve (AUC) is threshold independent, widely used to evaluate the predicted accuracy of the model performance (Phillips et al., 2004). The model performance was classified as poor (0.5-0.6), fair (0.6-0.7), good (0.7-0.8), very good (0.8-0.9), or excellent (0.9-1) and higher AUC values indicate better model performance (Swets, 1988). Partial AUC was calculated using ntBox (Lira-Noriega et al., 2020). MaxEnt output map was converted into binary ‘presence–absence’ map using 10 percentile training presence cloglog value to define suitable and unsuitable area for species. This threshold selects the value above which 90% of the training locations are correctly classified and is one of the most common thresholds used in MaxEnt habitat suitability models (Zarzo-Arias et al., 2019). All GIS analysis was performed in QGIS Ver 3.18.

### Niche shift analysis

To test for niche overlap between different species of *Pila* we performed Principal Component Analysis (PCA-env) approach as proposed by Broennimann et al., (2012). PCA method transforms environmental variables into two-dimensional space projected onto 100×100 PCA grids of cells bounded by minimum and maximum PCA values of background data. Niche overlap is measured using Schoener’s D metric and was used to measure the niche overlap, which varies from 0 (no overlap) to 1 (complete overlap). Niche equivalency and similarity are statistically tested from the density in environmental space (Broennimann et al., 2012). The niche equivalency test was used to assess whether the niches of two entities being compared are equal (display constant overlap), moderately similar (show some overlap) or significantly different (display no overlap) when occurrences are randomly shuffled across the ranges. The equivalency was performed by comparing the overlap (D) for native and introduced ranges with a null distribution. If the observed niche values were lower than the overlap value from the null distribution (P < 0.05), then the null hypothesis of niche equivalency is rejected (Broennimann et al., 2012). The niche equivalency test is limited by the fact that evaluation is not accounting for environmental conditions of the surrounding space of available habitat (Warren et al., 2008). Hence, the niche similarity test was adopted to assess whether the niches of two entities are more similar or different than would be expected by chance (Warren et al., 2008). This analysis was carried out using the “Ecospat” package (Di Cola et al., 2016) in R (R core team 2018) with 100 replicates to ensure that the null hypothesis can be rejected with high confidence. Further, this approach is considered robust as it uses kernel density smoothing to mitigate the effects of sampling bias (Petitpierre et al., 2012).

### Mantel test

In order to determine whether the speciation events can be a function of isolation by distance, mantel tests were performed separately for the two species complexes. The pairwise genetic distance between all samples were obtained from Mega7 under kimura 2 parameter model (Table S5). Next, the correlation between the geographic and genetic distance were measured using mantel test. The analyses were performed in R using the package Ecodist. Furthermore, we explored whether climatic factors play any role in governing the said speciation events. The values of each of the 19 bioclimatic variables for each sampling point were extracted from QGIS Ver 3.18. Thereafter, correlation between these variables were calculated in Excel and correlated variables were excluded from further analysis (Table S5). We carried out a partial mantel test in R using the package ecodist, in order to find out the correlation between genetic distance and the bioclimatic variables removing the effects of geographic distance. The analyses were carried out separately for *P. globosa* and *P. virens* complex.

### Analysis of genetic structure

We wanted to ascertain if water divides separating one river basin from another act as barriers to gene flow. Hence presence of genetic structure within the species complexes were investigated using BAPS in a Bayesian framework. BAPS Since the analyses were based on mitochondrial genes, hence, the ‘clustering with linked loci’ option was selected. In the absence of any prior information, the maximum number of clusters were fixed as ten and five respectively and analysed independently.

### Divergence dating

Divergence dating was carried out in Beast v1.8.2 (Drummond et al., 2012). The calibrations, and other prior parameters were specified in accordance with Sil et al., 2020. Two additional individuals were included in the analyses: one individual from clade C and one individual from clade D. The analyses were run for 100 million generations with a sampling frequency of 1000. The resultant topology was slightly different to the topology retrieved in Sil et al., (2020). Hence, another analysis was carried out where the topology was constrained to be similar to Sil et al., 2020. The marginal likelihood of both the analyses were estimated using path sampling and stepping stone sampling method. A total of 100 path steps were performed for 500,000 generations. The marginal likelihood values were compared using Bayes Factor test. The ESS values were examined in Tracer. The output of the analysis with significantly higher marginal likelihood was summarized in Treeannotator and visualized in Figtree.

## Results

### Phylogenetic and Species delimitation analyses

The *P. globosa* phylogeny consisted of four clades (Figure 1). All individuals from Gangetic basin formed a clade (A) that was sister to all other individuals. The individual collected from Orissa formed two sister clades (B and C respectively). All individuals from Northeast (Brahmaputra basin) were monophyletic (D) and sister to clade B and C. The single individual collected from Cachar was sister to all other individuals in the clade D. All the clades were well-supported. The species delimitation analyses showed that *P. globosa* is in actuality a species complex consisting of four cryptic species — each corresponding to the clades A, B, C, and D respectively. The clade D was identified as a previously described species from Cachar, Northeast India, called *P. olea*. From hereon clade D will be referred to as *P. olea*. However, as *P. globosa* is paraphyletic with respect to the former, the whole complex will be referred to as *P. globosa* complex.

**Figure 1:**
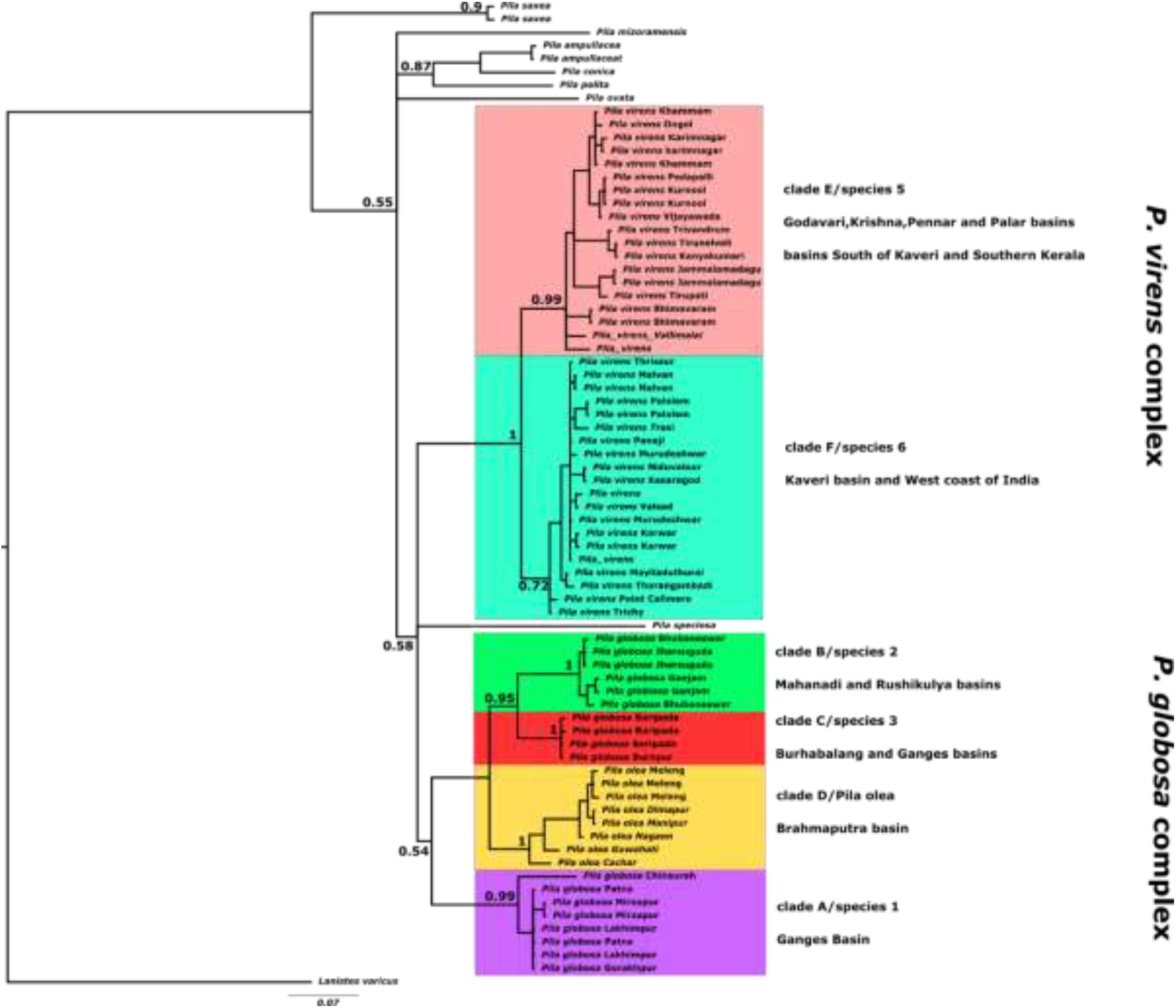
Bayesian inference phylogeny and species delimitation of *Pila*

The *P. virens* phylogeny was composed of two moderately supported clades (E and F respectively). Clade E consisted of all individuals from Andhra Pradesh and Telangana (Godavari, Krishna and Pennar river basins), and a few individuals from Southern Tamil Nadu, and one individual from the Southern part of Kerala, near Trivandrum. Clade F, on the other hand, consisted of all individuals from the west coast of India, from Kerala in the south, till southern Gujarat in the north. It also consisted of a few individuals from Kaveri basin of Tamil Nadu. Clade E and F were found to be two different species, as shown from the results of the species delimitation analyses. *Pila saxea* was retrieved to be one single species.

### Geometric morphometrics

The PCA plot of different putative species belonging to *P. globosa* and *P. virens* species complex (Figure S1) did not show any grouping based on morphometric characters suggesting that the shape of shell cannot be used as a deterministic feature to delimit species. Furthermore, none of the putative species showed significant distance in the morphological disparity analyses.

### Morphometric analyses

The PCA plot of the putative species from P. virens complex were overlapping but largely distinct. PC1 alone contributed 37% of the total variance (Figure S2). Likewise, the ANOVA showed significant difference between the two putative species (Table S4). Based on the silhouette width the final PAM analysis was run based on six clusters. However, the results showed that the three biggest clusters have representatives from both the species. The other three clusters consisted of only three individuals each. The putative species constituting P. globosa species complex showed more overlap in the PCA plot. No significant difference was found in the ANOVA analysis (Table S4). The PAM analysis retrieved two clusters, both constituting of individuals from all four putative species (figure S3).

### Nice modelling and niche divergence

The niche modelling results (Figure 2) shows that *P. saxea* and clade F from *Pila virens* complex have niches restricted to the Western Ghats and the West Coast respectively. Clade E from *Pila virens* complex is mostly restricted to part of Deccan peninsula and the East Coast of India. On the other hand, clade A from *P. globosa* complex has distribution mostly in the Gangetic plains, but extending to Western part of India and towards south-east region. Though *P. olea* is restricted to NE India, the ENM map predicts suitable habitats all along east coast and parts of Deccan peninsula. The most ambiguous is clade B of *P. globosa* complex. The ENM predicts most part of India is suitable. This is purely due to sampling artefact as we have only less than six records for this species.

**Figure 2:**
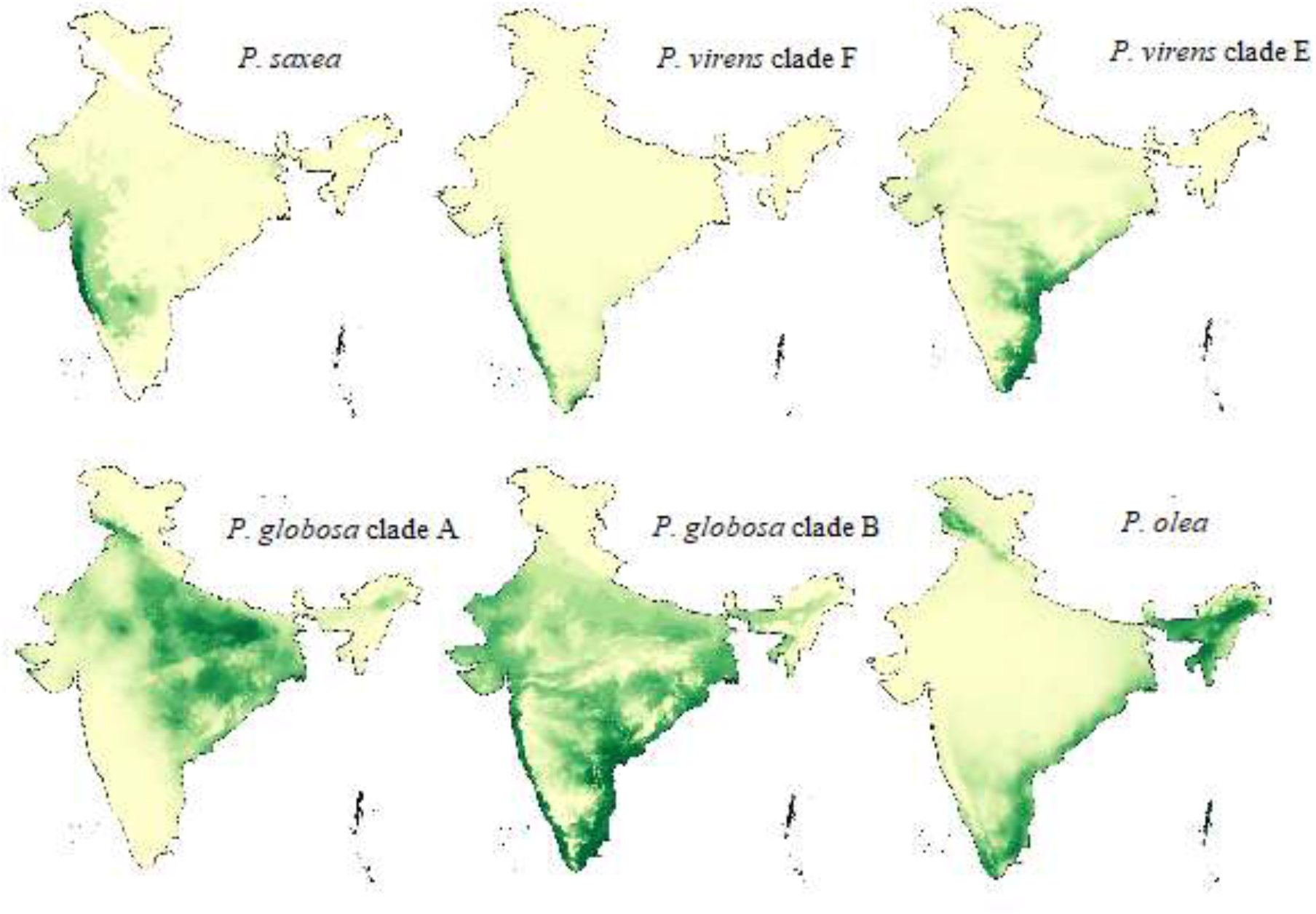
Predicted distribution of different described and putative species of Indian *Pila*

The niche overlap analysis (Figure S4) suggest that there is a partial overlap between Clade A and B from *P. globosa*., clade A (*P. globosa*) and clade E (*P. virens*) and P. olea. On the other hand, within *P. virens* complex, clade F has very small or insignificant overlap in the niche with clade E and partial with *P. saxea*. There is no overlap with other pairs of species. Incidentally, there is little overlap within *P. virens* complex.

### Mantel test

The genetic distance between the individuals from *P. globosa* and *P. virens* complex were positively correlated with the geographic distance. Some of the environmental variables were positively correlated as well. However, when the geographic distance was controlled for, only a few variables came out to be positively correlated with the genetic distance and significant (see table S5 for details).

### Analysis of genetic structure

No different were noticed between analyses using five and ten maximum number of clusters (Figure S5). Four clusters were recovered from *P. globosa* species complex (p, q, r and s), each corresponding to the putative species retrieved from the mPTP analysis (A, B, C and *P.olea*). Two of the clusters (p and q) were restricted to Ganges and Brahmaputra basin respectively. The third cluster (r) was shared between Mahanadi and Rushikulya basin. The fourth cluster (s) was shared between Burhabalang and Ganges basin. The *P. virens* complex gave rise to 4 clusters as well, three of them (w, x and y) belonging to putative species E, and the last one to putative species F. Out of these the cluster w corresponds to all individuals collected from Krishna basin and Govavari, and one individual from Pollar basin. The individuals sampled from Pennar basin constituted the cluster x. The third cluster y was composed of individuals from basins south of Kaveri and west flowing river m at the southern end of west coast of India. Although, putative species F is disjunctly distributed in Kaveri basin in the east and all through the west coast in the west, all individuals formed a single cluster, z.

### Divergence dating

The results of the divergence dating analysis was similar to Sil et al. (2020) (Figure 3). However, P. virens complex was retrieved as sister to P. globosa complex whereas it was retrieved as sister to all other Asian Pila in Sil et al 2020. The Bayes Factor test suggested that the Sil et al., (2020) topology was significantly better than that of the current topology (log BF 15.26 from path sampling and 15.05 from stepping stone sampling analyses respectively). Hence, we are going to discuss the results only from the constraints topology. Clade B, C, and *P. olea* separated from clade A (Ganges basin), belonging to *P. globosa* complex during Oligocene—Miocene (27.3—11.3 million years ago (mya)). Divergence between *P. olea* (Brahmaputra basin) and B-C took place around 17.1—6.0 mya during Miocene. Finally clade B (Mahanadi basin) diverged from C around 10.4—2.6 mya (Miocene—Pliocene). The two putative species from *P. virens* complex (clade E and F) separated from one another during Miocene epoch. (19.2—5.9 mya). The Southern Peninsular Indian population of clade E split off from the Godavari-Krisshna-Pennar basin population during Pio—Pleistoceen (4.6—0.5 mya). In clade F the West coast population diverged from the Kaveri basin population around the same time (3.3—0.3 mya).

**Figure 3:**
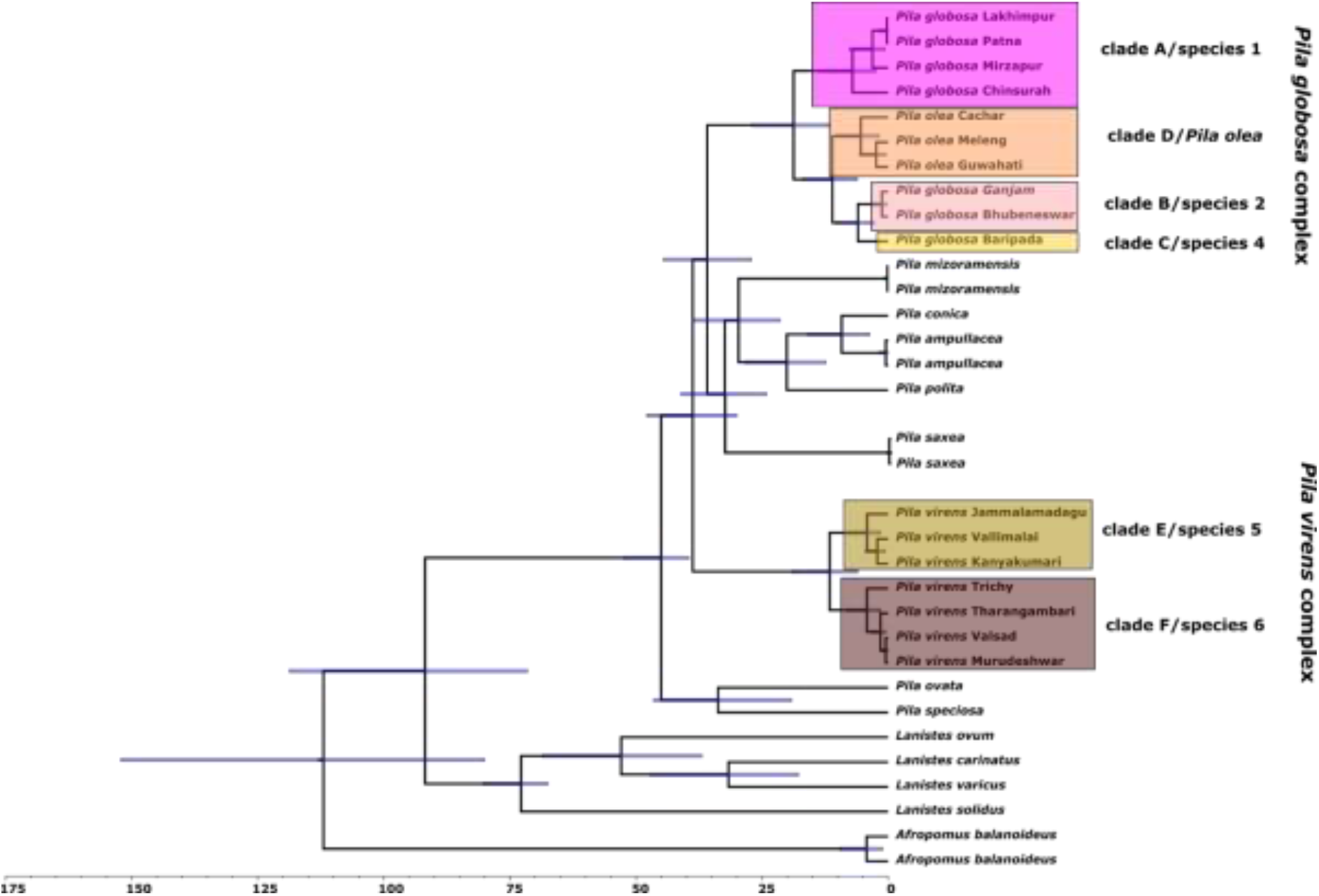
Divergence dating analysis

## Discussion

The manuscript aimed to understand the cryptic diversity in the genus *Pila* and the factors governing the diversity. The molecular data from the COI gene indicate that these species complexes constitute of several putative species. The environmental data garnered support in favour of these putative species, whereas, the morphometric data failed to distinguish between them. In the next few sections, we discuss the implications of such a pattern, as well as the ecological factors governing such diversity.

### Environmental and genetic segregation in the absence of morphological divergence

Both *P. globosa* and *P. virens* exhibited hitherto undetected cryptic diversity. *Pila globosa* complex constituted of three cryptic species and a previously described species *P. olea*. *P. virens* complex was composed of two cryptic species, according to the COI based species delimitation analyses, mPTP. The ecological niche model as well as niche overlap analysis further showed that the projected distribution ranges of all the putative species are less overlapping with each other. However, the morphometric data failed to show any differences between the putative species. There can be a few possible explanations to this pattern. Ongoing or past gene flow can limit the degree of morphological divergence (Moore et al., 2007). However, this seems unlikely, because, the ranges of the putative species do not overlap. More insight could have been gathered from the topological differences between the mitochondrial and nuclear trees. However, the nuclear concatenated tree was unresolved, hence, uninformative. Secondly, these putative species overall occupy similar lacustrine habitats. Hence, niche conservatism might have promoted absence of strong divergent selection (Fišer et al., 2018). The absence of divergent selection might have prevented morphological divergence. Lastly, it has shown that shell morphology in a population is often governed predominantly by ecological factors, such as predator pressure, and exhibit phenotypic plasticity (Brönmark et al., 2011; Trussell, 2000). Hence, a large variation may exist within these putative species, which might overshadow the interspecies differences. The differences in shell morphology if they exist, might be qualitative, thus not captured by morphometric analyses. Other characters such as anatomical characters have to be examined in order to find interspecies differences.

### Allopatric speciation between putative species

No two different putative species were collected from the same sampling location. Furthermore, individuals from geographically close locations chiefly consisted of members of the same putative species. The results from mantel test indicated at presence of geographic structure amongst both *P. globosa* and *P. virens* species complexes. Similar geographic structure was observed in the results from the BAPS analyses as well. The niche modelling analyses showed that the putative species show little or no overlap in their geographic range. All these evidences suggest that the mode of speciation was allopatric not sympatric parapatric. Although in cases observed allopatry can be secondary, but presence of strong geographic structure deems it unlikely. One has to investigate signatures of historical gene flow in order to ascertain whether it was parapatric or allopatric (Salgado-Roa et al., 2021). In the next section we discuss the various potential drivers behind these speciation events.

### Factors governing speciation

Allopatric speciation can result from various factors, such as isolation by distance, geographic (Chan et al., 2018; Mayr, 1954) or climatic barriers (Tomasello et al., 2020), adaptation to different climatic conditions (Jezkova and Wiens, 2018). Isolation by distance seems to be a major factor in driving speciation in the focal taxa. This is expected owing to the low dispersal rate of freshwater snails. The genetic distance is also correlated to various climatic factors, suggesting that they also might have played a role in driving speciation in these groups. However, in case of *P. globosa* complex, climatic factors does not seem to have a large effect on genetic distance when effects of the geographic distance was removed. It is likely that the existing differences in climatic niche between species arose as a result of their geographic separation. In case of the *P. virens* complex however, four climatic variables, namely, BIO1, BIO 4, BIO6 and BIO 12 seemed to have significantly affected the genetic distance. This is plausible, since, putative species F are largely distributed in the west coast of India which enjoys higher rainfall than the distribution range of putative species E. Such divergence along the climatic niche axes can give rise to locally adapted populations and eventually to speciation (Linck et al., n.d., Keller and Seehausen, 2012). Moreover, the split between species E and F, span much of Miocene and overlaps with the establishment of the Indian monsoon and orogenesis in the Himalayas and Tibetan plateau. These events are largely responsible for the advent of seasonality and differences in rainfall patterns (Clift et al., 2008; Molnar and Rajagopalan, 2012; Nelson, 2007). This time frame was also associated with a series of diversification in wet-adapted and freshwater taxa (Agarwal et al., 2019; Agarwal and Karanth, 2015; Anoop et al., 2018; Sidharthan et al., 2020). Hence, the results suggest that climatic changes in the subcontinent might be at least partially responsible for speciation in *P. virens* complex. However, the adaptations to various climatic niche could be secondary as well. Moreover, we need to detect the specific adaptations these species underwent in response to climatic niche divergence, as well as how those changes influenced reproductive isolation to obtain a better understanding of this possible ecological speciation event (Nossil, 2012).

Interestingly, most putative species of *P. globosa* complex seem to be restricted to certain river basins. Such patterns have been observed in freshwater snails and other freshwater taxa (Reid et al., 2013). To illustrate, individuals from clade A and *P. olea* were restricted solely to Ganges and Brahmaputra basins, respectively. Individuals of clade B were predominantly collected from Mahanadi basin. However, they were also distributed in Rushikulya basin. Rushikulya basin is adjacent to Mahanadi basin. Hence, several possible events could have led to this. A river capture event in the past might have led to sharing of species between basins. Other possibilities include dispersal through a canal, or dispersal during flooding, or avian mediated dispersal. If the water-divide between basins is not very high then, *Pila*, being an amphibious species could also have dispersed over land. However, since, we see strong geographic structure the last two possibilities do not seem likely. Individuals of clade C were collected from two locations, from Budhabalanga and Ganges basin respectively. This can also result from the reasons listed above. Ganges basin is separated from the basins in Odisha by the Vindhyas and Chhotanagpur plateau and from Brahmaputra sub-basin by Brahaputra ridge. River capture events can facilitate cross-basin dispersal (Anoop et al., 2018; Sidharthan et al., 2020). However, no information about any river capture events exists between these basins. One known paleohydrological event that could have shaped the species in *P. globosa* complex, is repeated marine incursion in the Bengal basin (Banerji, 1984), which might have separated the populations allopatrically, which eventually led to speciation. Conversely, rare flooding events also could have led to inter-basin dispersal. During mid-Miocene higher rainfall might have facilitated inter-basin connections (Sidharthan et al., 2020).

None of the putative species in the *P. virens* complex were restricted to a single river basin. Clade E is disjuctly distributed in two areas. The northern group is distributed from Godavari basin in the north to Pollar basin in the south. It has not been found in Kaveri basin. It again reappears south of the Kaveri basin, also near Tivandrum on the Southern part of West coast. Individuals from putative species F were found in parts of Kaveri basin and all over the west coast, starting from north of Trivandrum to Southern Gujarat. It is tempting to explain this pattern as a result of hydrological divide between the west and east flowing rivers in Peninsular India. However, the entire west coast clade is nested within the Kaveri basin clade and seems to have resulted from a recent expansion. Repeated marine incursions in the west coast might have extirpated previous populations if any. The split between putative species E and F might have taken place between Kaveri basin and other peninsular Indian basins north of Kaveri. The Mulki-Pulikat lake (MPL) axis, a ridge spreading from west coast to the east coast of India, just north of Kaveri basic (Subrahmanya, 1995), might have acted as a barrier to dispersal. The split between clade E and F dates back to Miocene. Much higher rainfall during mid-Miocene might have facilitated dispersal between drainage basins. Conversely, a river capture event between Kaveri and basins north of the MPL axis also could have led to this. A section of the Kaveri basin, between Shivasamudram and Vellore underwent tectonic rejuvenation during late Miocene or post-late Miocene (Jain et al., 2020; Kale et al., 2014; Radhakrishna, 1993). The upper Kaveri river initially used follow the course of the Pollar river, which flows north of the present day Kaveri river, while the lower Kaveri river used to flow from near Palghat along its usual course through the plains of Tamil Nadu. The upliftment caused a series of river capture events, that joined the upper and lower Kaveri river to form its current channel. Given that clade E is still found in Pollar basin, it is likely that river capture event caused the split between clade E and F. The divergence time-frame of the two clades is also concurrent with the event. The Southern Peninsular Indian population of putative species E diverged recently from the northern population. The dispersal to the Southern part might have taken place during the Pleistocene sea level drops, when part of the continental shelf was exposed.

River basins are known to drive speciation and genetic structure in freshwater species. The degree to which the evolutionary trajectory is shaped by the river basins is of course contingent upon the biology of the species. Not many studies have explored how genetic diversity is shaped by hydrological systems in freshwater snails. Much of the studies targeted lotic systems. It was intriguing to observe that the evolutionary processes in generalist groups such as *Pila* was moderately shaped by river basins.

### Conclusion

The results show that both *P. globosa* and *P. virens* consists of multiple putative species. These species are morphologically conserved but genetically and ecologically distinct. The speciation event in these two groups took place during Miocene. The diversification in these groups is closely tied to fact that freshwater organisms seldom disperse across river basins. The humid climate that prevailed during mid-Miocene might have facilitated cross-basin dispersal and the subsequent aridification during late Miocene might have isolated the populations. Alternately cross-basin dispersal could have been facilitated by river capture events, and the isolated by marine transgression events. Some of the population level dispersals might have been facilitated by periods of humid climate during the Pleistocene. Similar studies of freshwater systems in the subcontinent will help shed more light on the trends of diversity and speciation in freshwater organisms in the subcontinent.

## Acknowledgments

The major part of the project was funded by grants from Dept. of Biotechnology, Govt. of India (BT/01/17/NE/TAX) to NAA. The part of the fieldwork was supported by Rufford Small Grant for Nature Conservation (19805-1) to MS and DST (Department of Science and Technology, Govt. of India)-SERB grant (EMR/2017/001213) to PK. Authors would like to thank Pruthviraj, Harshal Bhosale, Tarun Singh, Aparna Lajmi, Chinta Sidhharthan, Kunal Arekar, Ananya Jana, Surya Narayan, Suhel Seikh, Shantanu Kundu for help during sample collection. Reshma Basak and Nipu Kumar Das provided important inputs towards geometric morphometry and morphometric data collection and analysis.

## Notes

### Competing Interest Statement

The authors have declared no competing interest.

