## Supporting Information for "Cryptic speciation in freshwaters: are speciation in lentic species shaped by paleohydrological events?"

Table S1. Details of samples used in the study

| Species | Clade | Basin | Location |
|---|---|---|---|
| *P. globosa* | A | Ganges | Chinsurah |
| *P. globosa* | A | Ganges | Patna |
| *P. globosa* | A | Ganges | Patna |
| *P. globosa* | A | Ganges | Mirzapur |
| *P. globosa* | A | Ganges | Mirzapur |
| *P. globosa* | A | Ganges | Lakhimpur |
| *P. globosa* | A | Ganges | Lakhimpur |
| *P. globosa* | A | Ganges | Gorakhpur |
| *P. globosa* | C | Budhabalanga | Baripada |
| *P. globosa* | C | Budhabalanga | Baripada |
| *P. globosa* | C | Budhabalanga | Baripada |
| *P. globosa* | C | Ganges | Burnpur |
| *P. globosa* | B | Mahanadi | Sambalpur |
| *P. globosa* | B | Mahanadi | Sambalpur |
| *P. globosa* | B | Mahanadi | Bhubaneswar |
| *P. globosa* | B | Mahanadi | Bhubaneswar |
| *P. globosa* | B | Rushikulya | Ganjam |
| *P. globosa* | B | Rushikulya | Ganjam |
| *P. globosa* | D | Brahmaputra | Guwahati |
| *P. globosa* | D | Brahmaputra | Meleng |
| *P. globosa* | D | Bramhapura | Meleng |
| *P. globosa* | D | Bramhaputra | Meleng |
| *P. globosa* | D | Bramhaputra | Dimapur |
| *P. globosa* | D | Bramhaputra | Sugandhi |
| *P. globosa* | D | Bramhaputra | Cachar |
| *P. globosa* | D | | Manipur |
| *P. virens* | E | | Vallimalai |
| *P. virens* | E | | Tirupathi |
| *P. virens* | E | Pennar | Jammalamadagu |
| *P. virens* | E | Pennar | Jammalamadagu |
| *P. virens* | E | Krishna | Kurnool |
| *P. virens* | E | Krishna | Kurnool |
| *P. virens* | E | Krishna | Ongole |
| *P. virens* | E | Krishna | Khammam |
| *P. virens* | E | Krishna | Khammam |
| *P. virens* | E | Krishna | Vijayawada |
| *P. virens* | E | Godavari | Pedapalli |
| *P. virens* | E | Godavari | Karimnagar |
| *P. virens* | E | Godavari | Karimnagar |
| *P. virens* | E | Godavari | Bhimavaram |
| *P. virens* | E | Godavari | Bhimavaram |
| *P. virens* | E | | Kanyakumari |
| *P. virens* | E | | Tirunelvelli |

| | | | |
|---|---|---|---|
| *P. virens* | E | | Trivandrum |
| *P. virens* | F | Kaveri | Trichy |
| *P. virens* | F | Kaveri | Point Calimere |
| *P. virens* | F | Kaveri | Mayladuturai |
| *P. virens* | F | Kaveri | Tarangampadi |
| *P. virens* | F | West flowing river | Trissure |
| *P. virens* | F | West flowing river | Niduvallur |
| *P. virens* | F | West flowing river | Kasadgod |
| *P. virens* | F | West flowing river | Trassi |
| *P. virens* | F | West flowing river | Palolem |
| *P. virens* | F | West flowing river | Palolem |
| *P. virens* | F | West flowing river | Murudeswar |
| *P. virens* | F | West flowing river | Murudeswar |
| *P. virens* | F | West flowing river | Karwar |
| *P. virens* | F | West flowing river | Karwar |
| *P. virens* | F | West flowing river | Panjim |
| *P. virens* | F | West flowing river | Malvan |
| *P. virens* | F | West flowing river | Malvan |
| *P. virens* | F | West flowing river | Valsad |

Table S2. Details of primers

| Gene name | Abbreviation | Length (bp) | Models | Primers | **References** |
|---|---|---|---|---|---|
| Cytochrome c oxidase subunit 1 | COI | 561 | Codon position 1 TrNef+G, codon position 2 HKY+I, codon position 3 HKY +G | LCO and HCO | (Folmer et al., 1994; Williams et al., 2003) |

| | | | | | |
|---|---|---|---|---|---|
| 18S ribosomal RNA | 18S rRNA | 465 | K80+G | 18SYLMFOR 18SYLMREV | (Stothard et al., 2000) |
| Histone H3 | Histone H3 | 279 | Codon position 1 and 2 K80+I, codon position 3 K80+I | H3F and H3R | (Colgan et al., 1998) |

Table S3. Details of the morphological variables used in the study

| Raw measurements taken | | | | | | | | | |
|---|---|---|---|---|---|---|---|---|---|
| Width (W) | Aperture height (AH) | Aperture width (AW) | Body whorl height (BWH) | Body whorl width (BWW) | Height of penultimate whorl (HPW) | Width of penultimate whorl (WPW) | Height of ante-penultimate whorl (HAW) | Width of ante-penultimate whorl (WAW) | |
| Ratios used for final analyses | | | | | | | | | |
| HAW/W | AH/HAW | AW/HAW | BWH/HAW | BWW/HAW | HPW/HAW | AW/AH | AW/BWW | BWW/BWH | WAW/WPW |
| P. globosa | | | | | | | | | |
| HAW/W | AH/HAW | | BWH/HAW | | HPW/HAW | AW/AH | AW/BWW | BWW/BWH | WAW/WPW |
| P. virens | | | | | | | | | |
| HAW/W | AH/HAW | BWH/HAW | HPW/HAW | AW/AH | | AW/BWW | BWW/BWH | WAW/WPW | |

Tables S4. Details of ANOVA

| P. globosa complex | Aperture Height/Ante-Penultimate height | Body whorl width/ Ante-Penultimate height |
|---|---|---|
| P value | 0.95 | 0.08 |
| P. virens complex | Aperture width/Body whorl width | Body whorl width/Body whorl Height |
| P | **0.03** | **0.01** |

Table S5. Details of relevant climatic variables used in mantel test

| **P. globosa species complex** | | | | | | | | | | |
|---|---|---|---|---|---|---|---|---|---|---|
| | **Geographic Distance** | | | | | | | | | |
| **Mantel r** | 0.59 | | | | | | | | | |
| **P value** | 0.001 | | | | | | | | | |
| | **BIO1** | **BIO2** | **BIO3** | **BIO5** | **BI08** | **BIO9** | **BIO12** | **BIO13** | **BIO14** | **BIO19** |

| | | | | | | | | | | |
|---|---|---|---|---|---|---|---|---|---|---|
| **Without controlling for distance** | | | | | | | | | | |
| **Mantel r** | 0.27 | 0.24 | 0.33 | 0.34 | 0.1 | 0.35 | 0.18 | -0.02 | 0.18 | 0.06 |
| **P value** | 0.002 | 0.002 | 0.001 | 0.001 | 0.035 | 0.001 | 0.007 | 0.63 | 0.01 | 0.15 |
| **After controlling for distance** | | | | | | | | | | |
| **Mantel r** | -0.08 | 0.07 | 0.23 | 0.023 | 0.06 | 0.09 | -0.04 | -0.08 | -0.17 | -0.21 |
| **P value** | 0.86 | 0.18 | 0.004 | 0.29 | 0.28 | 0.093 | 0.7 | 0.8 | 1.0 | 1.0 |
| **P. virens species complex** | | | | | | | | | | |
| | **Geographic Distance** | | | | | | | | | |
| **Mantel r** | 0.36 | | | | | | | | | |
| **P value** | 0.001 | | | | | | | | | |
| **Without controlling for distance** | | | | | | | | | | |
| | **BIO1** | **BIO4** | **BIO6** | **BIO12** | **BIO14** | **BIO18** | **BIO19** | | | |
| **Mantel r** | 0.34 | 0.52 | 0.31 | 0.48 | 0.12 | -0.03 | 0.03 | | | |
| **P value** | 0.001 | 0.001 | 0.001 | 0.001 | 0.019 | 0.7 | 0.21 | | | |
| **After controlling for distance** | | | | | | | | | | |
| **Mantel r** | 0.23 | 0.42 | 0.19 | 0.44 | 0.003 | -0.08 | -0.08 | | | |
| **P value** | 0.002 | 0.001 | 0.003 | 0.001 | 0.5 | 0.9 | 0.9 | | | |

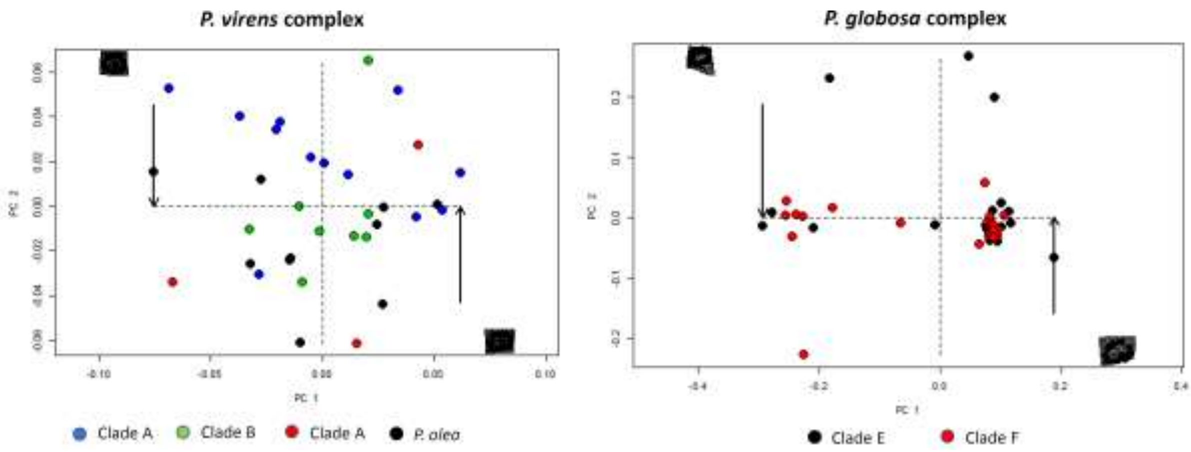


Figure S1: Geometric morphometrics of *Pila*

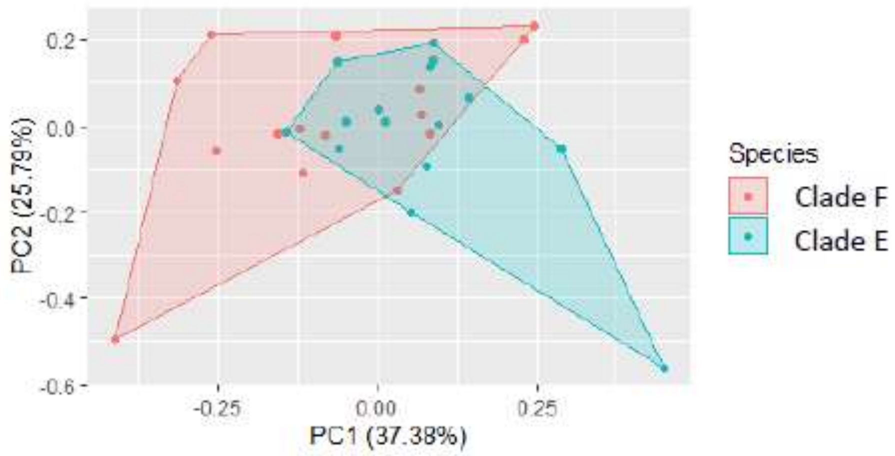


***P. virens* complex**

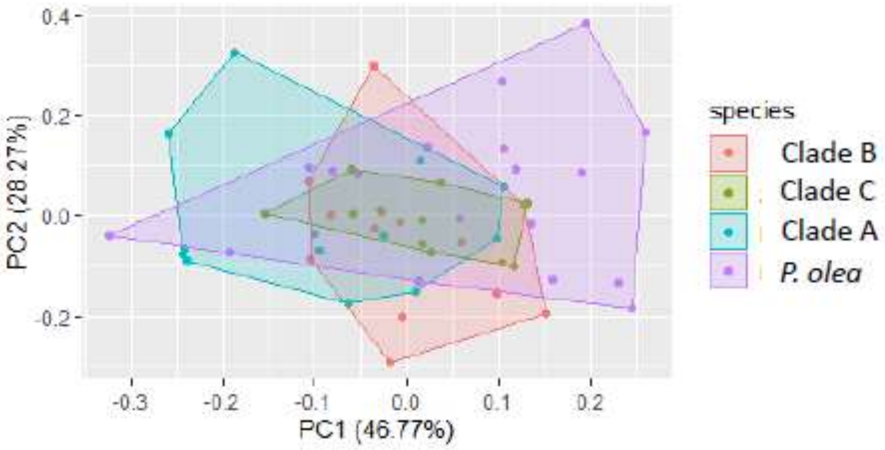


***P. globosa* complex**

Figure S2: PCA analysis of morphometric variables of *Pila*

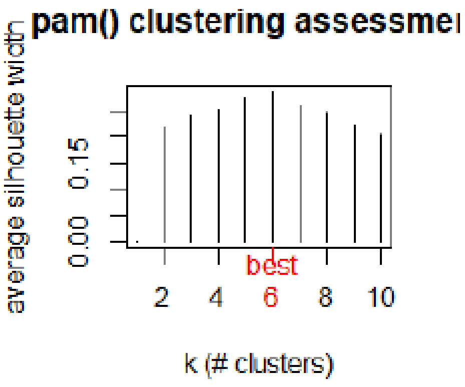


***P. virens* complex**

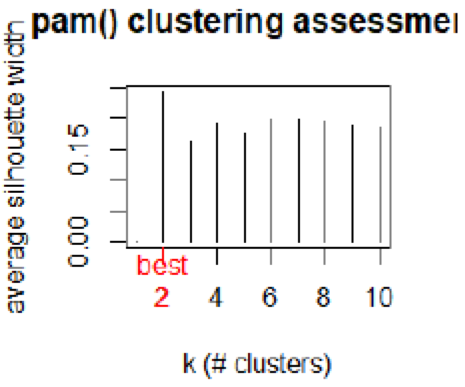


***P. globosa* complex**

S3: PAM analysis of morphometric variables of *Pila*

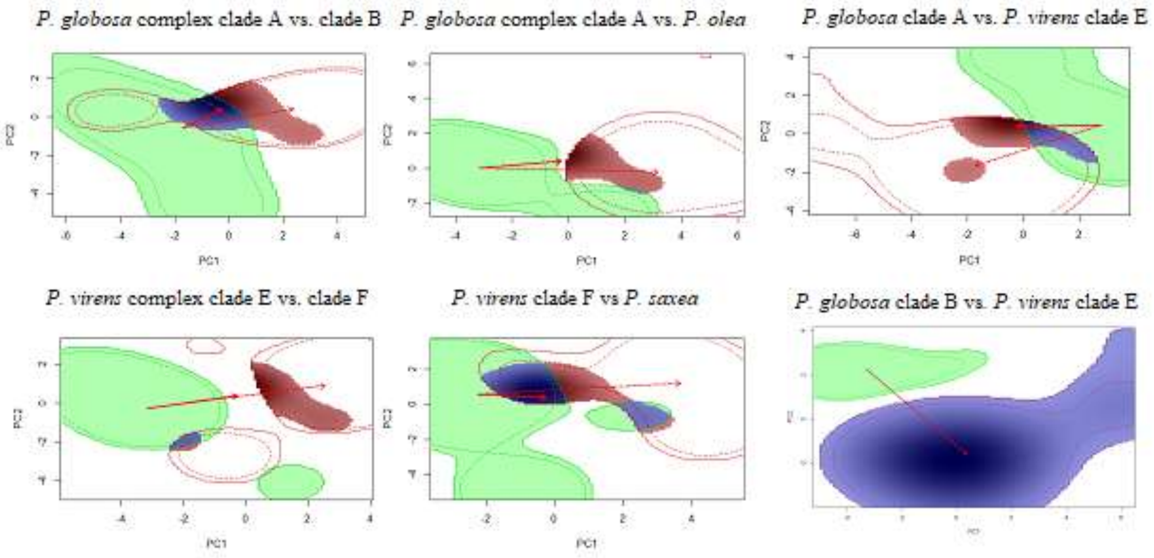


Figure S4: Niche overlap analysis of *Pila*

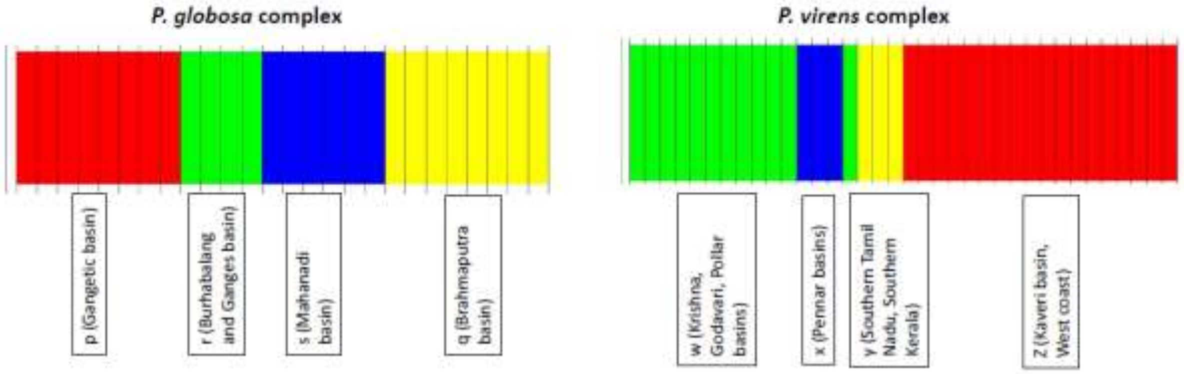


Figure S5: analyses of genetic structure.
